# Pathogens in exacerbations of chronic rhinosinusitis differ in sensitivity to silver nanoparticles

**DOI:** 10.1101/2022.01.03.474872

**Authors:** Joanna Szaleniec, Agnieszka Gibała, Joanna Stalińska, Magdalena Oćwieja, Paulina Żeliszewska, Justyna Drukała, Maciej Szaleniec, Tomasz Gosiewski

## Abstract

**Purpose:** The significance of the microbiome in chronic rhinosinusitis (CRS) is not clear. Antimicrobials are recommended in acute exacerbations of the disease (AECRS). Increasing rates of antibiotic resistance stimulate research on alternative therapeutic options including silver nanoparticles (AgNPs), sometimes referred to as “colloidal silver”. However, there are concerns regarding the safety of silver administration and the emergence of silver resistance. The aim of this cross-sectional observational study was to assess the sensitivity of sinonasal pathogens to AgNPs and compare it with the toxicity of AgNPs for nasal epithelial cells.

**Methods:** Negatively charged AgNPs (12±5 nm) were synthetized using tannic acid. The minimal inhibitory concentration (MIC) for pathogens isolated from patients with AECRS was approximated. Cytotoxicity of AgNPs was tested *in vitro* on human nasal epithelial cells line.

**Results:** 48 clinical isolates and 4 reference strains were included in the study (*Staphylococcus aureus, Pseudomonas aeruginosa, Escherichia coli, Klebsiella pneumoniae, Klebsiella oxytoca, Acinetobacter baumanii, Serratia marcescens, Enterobacter cloacae*). The MIC values differed between isolates, even within the same species. All the isolates (including antibiotic resistant) were sensitive to AgNPs in concentrations nontoxic to human cells during 24 h exposition. However, 48 h exposition to AgNPs increased toxicity to human cells, narrowing their therapeutic window and enabling 19% of pathogens to resist the AgNPs’ biocidal action.

**Conclusions:** AgNPs are effective against most pathogens isolated from patients with AECRS, but sensitivity testing may be necessary before application. Results of sensitivity testing for reference strains cannot be extrapolated to other strains of the same species.

## Introduction

Chronic rhinosinusitis (CRS) is an inflammatory disease of the sinonasal mucosa. Understanding of the complex relationships between the inflammatory process and the microbiota of the sinuses is still in its infancy. The role of bacteria seems to be more evident in acute exacerbations of chronic rhinosinusitis (AECRS). If sudden worsening of symptoms is accompanied by purulence in the sinuses, the incident is attributed to bacterial infection and antibiotic treatment is recommended [1, 2]. Increasing antibiotic resistance of sinonasal pathogens is one of the factors that limit treatment efficacy in AECRS [3]. As a result, there is growing interest in novel therapeutic options.

Intranasal preparations of silver nanoparticles (AgNPs) are proposed as an alternative treatment for sinonasal infections if antibiotic therapy is ineffective. Silver nanoparticles (AgNPs) are objects of any shape and dimensions in the range from 10^−9^ m to 10^−7^ m built from silver atoms. Depending on the methods and stabilizing agents used to synthesize the AgNPs, they differ significantly in physicochemical and biological properties [4, 5]. The AgNPs dispersed in liquid media are sometimes referred to as “colloidal silver” [6]. However, this term is not precise and may lead to misunderstandings. Many over-the-counter preparations marketed as “colloidal silver” contain various unspecified forms of silver, but no well-characterized silver nanoparticles [7].

Potential medical applications of AgNPs evoke either enthusiasm or serious concerns. The antimicrobial properties of silver have been known for millennia. Silver has been shown to be effective against a wide variety of microbes, including Gram-positive and Gram-negative bacteria, yeast and fungi [8]. It damages many vital structures in the cells and therefore it was believed that resistance to silver was very unlikely to emerge [9, 10]. Unfortunately, recent studies prove that bacteria can develop manifold mechanisms of silver resistance, some of which additionally result in cross-resistance to antibiotics [11]. Research on AgNPs toxicity provides contradictory results. Some studies report acceptable safety of AgNP preparations [6, 12, 13], while others provide alarming data on cytotoxicity and accumulation in many organs including the brain after intranasal delivery [14]. The toxicity of silver preparations undoubtedly depends on dosing and formulation but reliable data on the optimal method of administration in sinonasal infections has not been established.

Recent studies show that bacteria in clinical settings sometimes harbour silver resistance genes and occasionally the clinical isolates can tolerate exceedingly high silver concentrations [15]. These findings undermine the conviction that silver is a universal antimicrobial and can be used for any infection without previous sensitivity testing. However, due to the lack of routine screening, the incidence of silver resistance in human infections is unknown.

The objective of our study was to assess if AgNPs could be effectively applied to fight pathogens isolated from patients with AECRS. The AgNPs in the study were obtained with the use of tannic acid. We tested the susceptibility to AgNPs in 48 pathogens isolated from 50 patients with AECRS. To our knowledge, to date, this is the largest survey on AgNP susceptibility in CRS patients [6]. In parallel experiments, we evaluated the toxicity of AgNPs for nasal epithelial cells *in vitro*. Our purpose was to determine if there is a potential therapeutic window, that is a concentration of AgNPs that is already effective against the pathogens but still safe for the host’s epithelium. The secondary goal was to evaluate the incidence of bacterial isolates that were resistant to AgNPs at concentrations non-toxic for human cells.

## Materials and methods

### Sample collection

The samples for this cross-sectional observational study were collected in the outpatient clinic of the Department of Otolaryngology of the University Hospital in Krakow in 2018 and 2019 (Jagiellonian University Medical College Bioethics Committee approval no. 1072.6120.208.2017). The eligibility criteria included: diagnosis of CRS according to EPOS 2012 diagnostic criteria [16], prior endoscopic sinus surgery (ESS), signs and symptoms of bacterial exacerbation [17] (worsening of symptoms, purulence in the sinonasal cavity), no antibiotic therapy for at least a week. Swabs were collected under endoscopic guidance from the pathological secretions from the middle nasal meatus or the infected sinuses. Bacterial isolates were stored at −80° C and thawed for experiments.

Reference strains were obtained from the American Type Culture Collection (*Escherichia coli* ATCC 25922, *Staphylococcus aureus* ATCC 29213, *Pseudomonas aeruginosa* ATCC 27857, *Klebsiella pneumoniae* ATCC 31488).

### Bacterial identification, antibiotic sensitivity testing and identification of antibiotic resistance mechanisms

The bacteria were inoculated on the Columbia blood agar with (OXOID) for Gram-positive aerobic cocci, on the chocolate base agar (OXOID) with bacitracin for *Haemophilus* and on the selective MacConkey agar (OXOID) for the isolation of Gram-negative bacilli. After 18-24 hours of incubation in the atmosphere containing 5% CO_2_ at 37° C the colonies were isolated. Subsequently, the bacteria were identified using a BD Phoenix (Becton Dickinson) automated microbiology system with appropriate test kits for Gram-negative and Gram-positive bacteria. *Haemophilus* rods were identified with discs containing factors V, X and bacitracin and on Müller–Hinton agar plates (OXOID) by incubating a McFarland 0.5 suspension with paper discs for 24 hours at 37 °C with access to CO_2_. Isolation and identification of bacteria and antibiotic susceptibility testing were performed as previously described [18] according to The European Committee on Antimicrobial Susceptibility Testing (EUCAST) 6.0[19]. Clinical breakpoints were interpreted according to EUCAST v. 8.0 [20]. For this purpose, standardized diagnostic discs containing a specific antibiotic were used. After bacterial growth was obtained, the zones of growth inhibition were measured using a calliper and the results were compared to EUCAST standards. The degree of antibiotic resistance and specific resistance mechanisms were derived from the diameters of the zones of inhibition according to EUCAST.

### Synthesis of silver nanoparticles

AgNPs were obtained using tannic acid as a reducing and stabilizing agent. In brief, 40 mL of 0.5 mM aqueous solution of tannic acid was introduced to 320 mL of 11 mM aqueous solution of silver nitrate. The reaction mixture was dynamically stirred on a magnetic stirrer at room temperature (*ca*. 25° C). Then, 30 μL of 25 wt % ammonia solution was introduced to the obtained reaction mixture. The stirring was continued for another 30 minutes. Finally, the obtained suspension was washed with MilliQ-water to remove unreacted reagents. The suspension was placed in an Amicon®filtration cell (model 8400) equipped with membranes made of regenerated cellulose with a nominal molecular weight limit of 100 kDa. The purification process was conducted under ambient conditions (25° C). The filtration cell was placed on a magnetic stirrer and gently mixed during the purification procedure. The progress in the purification was monitored via conductivity measurements where the conductivity was determined using a CPC-505 pH-meter/conductometer (Elmetron) equipped with a conductometric sensor EC-60 for every 30 ml of collected effluents. The purification process was conducted until the conductivity of the effluents stabilized at 20 μS cm^−1^ and the pH attained a value of 5.8 [5].

The concentration of AgNP in the suspension was determined with a densitometric method according to a previously established protocol [21]. Furthermore, the concentrations of Ag^+^ ions and AgNPs in the suspension were validated with ICP-OES (Perkin-Elmer OPTIMA 2100DV). After separation of AgNP from the solution by ultrafiltration (Amicon, 30 kDa) the respective fractions were dissolved in 70% HNO_3_ and subsequently in MiliQ-water before ICP-OES analysis. From each fraction, three independent samples were collected and analyzed in triplicate. The ICP-OES analysis validated the total concentration of AgNP established with the densitometric method (respectively 217±3 vs 214 mg L^−1^) and did not detect free Ag^+^ ions in the effluent within the limit of the detection method.

The optical properties of AgNP suspension were evaluated with the use of a UV-2600 spectrometer (Shimadzu). The morphology and size distribution of AgNPs were determined based on micrographs recorded using a JEOL JSM-7500F electron microscope working in the transmission mode (TEM). The obtained micrographs were analyzed with the use of MultiScan software (Computer scanning system). The histograms were generated from the analysis of no less than 500 AgNPs. The hydrodynamic diameter and zeta potential of AgNPs were determined from the measurements of diffusion coefficients (*D*) and electrophoretic mobility (*μ_e_*) which were conducted using a Zetasizer Nano ZS instrument (Malvern).

The AgNPs obtained using tannic acid were selected for this study due to their promising antibacterial properties observed in our preliminary experiments [5].

### Determination of the minimal inhibitory concentration (MIC) of the AgNPs

Evaluation of MIC in a liquid medium using turbidimetric measurements turned out to be impossible due to interference between the AgNPs at higher concentrations and the instrument’s readouts. To overcome this problem, an alternate method was developed, which approximates MIC according to the procedure presented in Fig. 1.

**Fig. 1.**
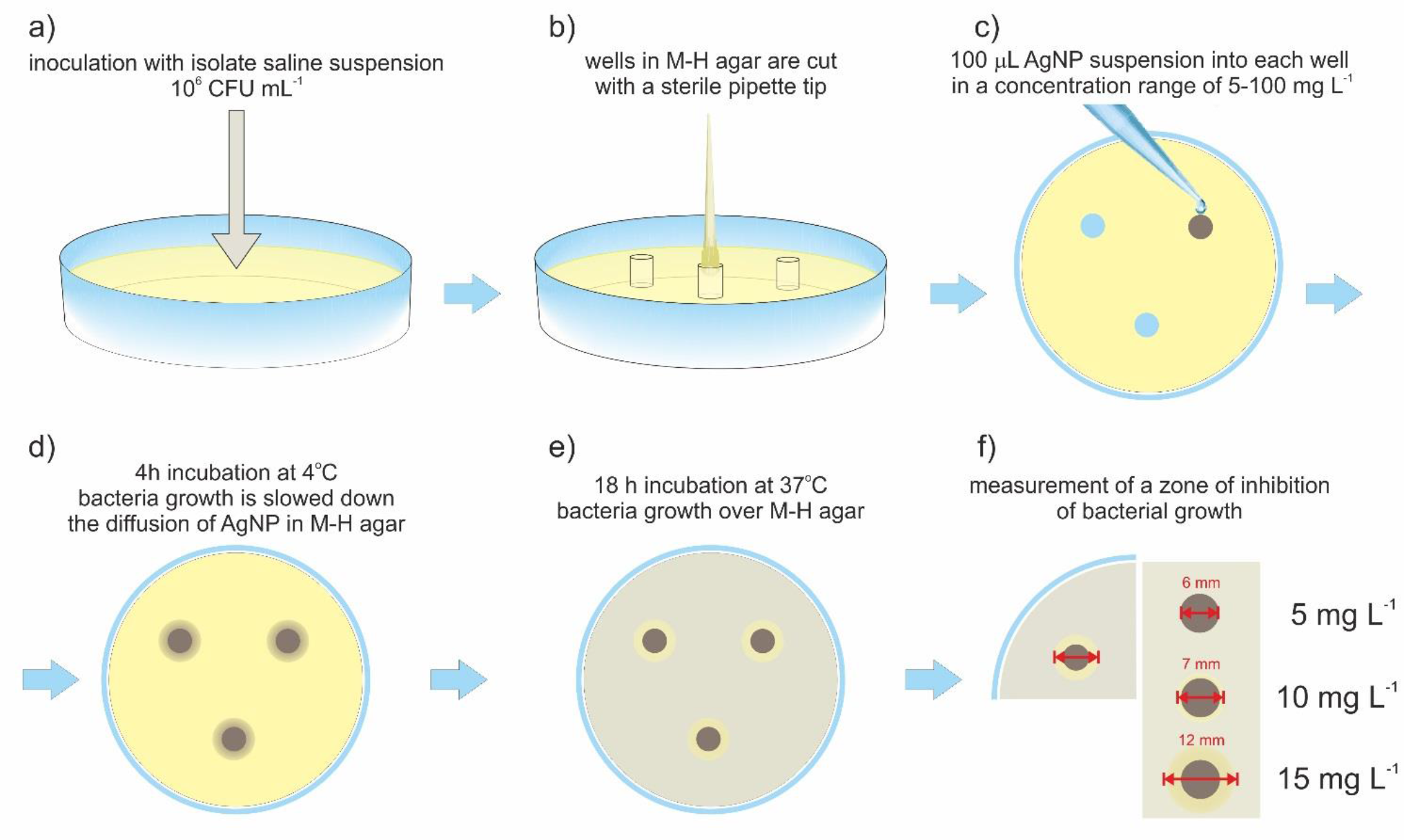
The method of MIC approximation used in the study. For the bacterial isolate presented in the figure, the actual MIC value is ≤ 10 mg L^−1^. M-H - Müller–Hinton agar.

To approximate the minimum inhibitory concentrations (MIC) of AgNPs, the suspension of the tested microorganisms 0.5 on the MacFarland optical density scale was diluted with saline 100 times to obtain a cell density of 10^6^ CFU mL^−1^ (colony forming unit) and inoculated on the surface of Müller–Hinton agar (OXOID) (Fig. 1a). Next, a sterilized pipette tip was used to form wells on the agar surface with a diameter suitable for a 200 μL pipette (Fig. 1b). Each well was filled with 100 μL of the AgNP suspensions in concentrations ranging from 5 to 100 mg L^−1^ (in a triplet for each concentration) (Fig. 1c). The concentration of AgNP suspensions was increased by 5 mg L^−1^ in subsequent wells. Pure medium without AgNPs was used as a control sample. Next, the plates were pre-incubation for 4 h at 4°C (Fig. 1d) to provide time for soaking the AgNP suspension into the agar gel and the proper diffusion of nanoparticles from the wells to the agar surface while arresting the growth of bacteria. After that time the plates were incubated at 37° C for 18 h (Fig. 1e). The zone of inhibition of bacterial growth around the wells was measured with Skjutmatt Digital calliper (Limit). All experiments were performed in triplicate.

Thus in this paper, MIC was defined as the lowest concentration of AgNPs in the well for which the zone of inhibition around the well was observed (e.g. 10 mg L^−1^ in Fig. 1f). In fact, the real concertation of AgNPs was inevitably diluted in the volume of the gel surrounding the well. Therefore, the value of the MIC based on the nominal concentration of the suspension introduced into the well should be treated as an upper limit of MIC. It means that the actual MIC value for each tested isolate was not higher than the lowest concentration of AgNPs initially introduced into the well around which the inhibition zone was observed. This method was not applicable for *Streptococcus pneumoniae* and *Haemophilus influenzae* because of the interactions between AgNPs and the growth media suitable for these species. Therefore, the results obtained for these bacteria were considered unreliable and were excluded from the analyses.

### *In vitro* primary cell culture and cell viability assays

Human nasal epithelial cells (HNEpC; PromoCell) were cultured in Airway Epithelial Cell Growth Medium (basal medium supplemented with BPE, EGF, insulin, hydrocortisone, epinephrine, triiodo-L-thyronine, transferrin and retinoic acid; PromoCell) at 37 °C in a 5% CO_2_ atmosphere. HNEpC cells at passage 5 were used in the experiments. Cells were plated in 48-well plates (BD Falcon) coated with type I collagen (Corning) at the initial density of 1 × 10^4^ cells/cm^2^, 24 hours prior to the experiments. The growth medium was then changed to the experimental cell culture medium containing nanoparticles or vehicle control (water). The experimental medium consisted of phenol red-free DMEM, prepared from powder using water or a mixture of water and 214 mg L^−1^ tannic acid-modified silver nanoparticles (AgNPs) suspension in water, and supplemented with 1 g L^−1^ glucose, NaHCO_3_ (Gibco), sodium pyruvate (Lonza), 1% bovine serum albumin (BSA), L-glutamine, and P/S (50 units/mL of penicillin and 50 μg/mL of streptomycin). Water (used as vehicle control) was a component of the cell culture medium, not an additive to the medium, and as such, the osmolality of the experimental medium was not compromised. A 48-hour incubation in the experimental medium had no negative effects on the viability of the cells compared to the PromoCell culture medium. Cell culture reagents were from Sigma unless otherwise specified.

#### 1.1.1 MTT assay

MTT assay was performed after the incubation in the presence of AgNPs, as previously described [22]. Following 2-hour incubation with MTT, formazan crystals were dissolved in 5 mM HCl in isopropanol and absorbance read at 540 nm. Experiments were performed in triplicate for each experimental condition. Data represent mean values expressed as a percentage of the vehicle control ± SD from three independent experiments.

#### 1.1.2 Trypan blue exclusion

After the 48-hour incubation in the presence of AgNPs, cells were harvested with StemPro Accutase (Gibco), centrifuged, and resuspended in DPBS. The cell suspension was diluted 1:1 with 0,4% trypan blue and counted in a hemocytometer [23].

### Statistical analysis

The normality of the distribution of the microbiological data was tested using the Kolmogorov-Smirnov test. The variables under consideration did not have a normal distribution, therefore non-parametric tests were used for further analyses. Mann-Whitney test was used for comparisons of two groups (e.g. comparisons of MIC values in Gram-positive and Gram-negative bacteria or bacteria with and without antibiotic resistance mechanisms) and the Kruskal-Wallis test was used to compare multiple groups (e.g. comparisons of MIC values between species). The tests were performed in Statistica 13.0. A p-value of < 0.05 was considered statistically significant. For the HNEpC viability assays, statistical analysis was performed using one-way ANOVA followed by Dunnett’s multiple comparisons test (GraphPad Prism 8.0).

## Results

Swabs from the sinonasal cavity were obtained from 50 patients with acute exacerbations of chronic rhinosinusitis (AECRS). The group included 27 women and 23 men aged 25-80 (mean age 51). All of the patients have undergone endoscopic sinus surgery in the past, so their sinuses were accessible for sampling. Most of the participants had chronic rhinosinusitis with nasal polyps (90%) and presented with comorbidities such as asthma (54%), aspirin-exacerbated respiratory disease (10%) or allergy (38%).

As presented in the flow chart in Fig. 2, the sensitivity to AgNPs was tested in 48 out of 97 isolates. Species considered nonpathogenic and bacteria that require media that inhibit the AgNPs were excluded.

**Fig. 2.**
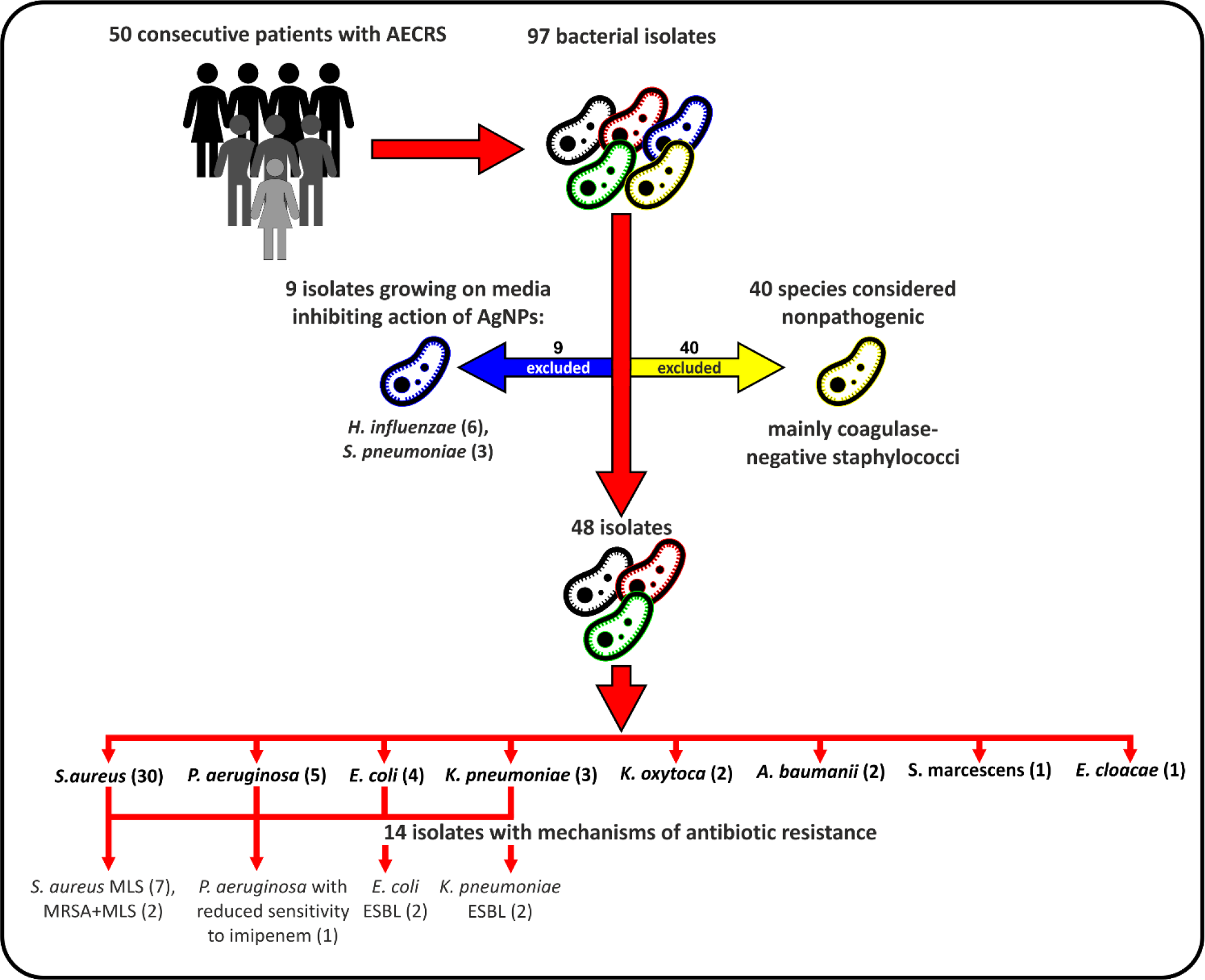
Flow chart detailing the selection of bacterial isolates for the study.

The AgNPs obtained with the use of tannic acid that were utilized in this study were quasi-spherical in shape, had a diffusion coefficient of 5.37 × 10^−7^ cm^2^s^−1^, hydrodynamic diameter 12±5 nm and zeta potential −52±3 mV (determined at the temperature of 37 °C and pH 5.8).

The upper limit of minimal inhibitory concentration (MIC) values of AgNPs for the clinical isolates and reference strains are presented in Table 1. Higher MIC values indicated lower sensitivity to AgNPs. Every experiment was performed in triplicate. The method of MIC determination was semiquantitative and provided results with an accuracy of up to 5 mg L^−1^. The MIC values determined by this method were highly reproducible. For every isolate, the same result was noted for all three replications (SD = 0). The MIC values for the clinical isolates ranged from 5 to 40 mg L^−1^ with a median value of 10 mg L^−1^. For each of the identified species, the sensitivity to AgNPs varied between isolates.

**Table 1.** The upper limit of minimal inhibitory concentration (MIC) of AgNPs for the reference strains and clinical isolates [mg L^−1^]. MIC values did not have a normal distribution, therefore median was used as a measure of central tendency and the interquartile range was used as a measure of statistical dispersion. Higher MIC values indicate lower sensitivity to the AgNPs.

| Species<br>(no of clinical isolates) | Reference<br>strain | Clinical isolates |  |  |  |
| --- | --- | --- | --- | --- | --- |
|  |  | median | min | max | interquartile<br>range |
| *Staphylococcus aureus* (30) | 5 | 10 | 5 | 40 | 15 |
| *Pseudomonas aeruginosa* (5) | 10 | 5 | 5 | 20 | 5 |
| *Escherichia coli* (4) | 5 | 25 | 10 | 25 | 7.5 |
| *Klebsiella pneumoniae* (3) | 5 | 25 | 10 | 30 | 20 |
| *Klebsiella oxytoca* (2) |  | - | 10 | 40 | - |
| *Acinetobacter baumannii* (2) |  | - | 10 | 10 | - |
| *Serratia marcescens* (1) |  | - | 25 | 25 | - |
| *Enterobacter cloacae* (1) |  | - | 30 | 30 | - |
| All clinical isolates (48) |  | 10 | 5 | 40 | 15 |
| All Gram-positive clinical isolates (30) |  | 10 | 5 | 40 | 15 |
| All Gram-negative clinical isolates (18) |  | 15 | 5 | 40 | 15 |

The lowest median MIC value was noted for *P. aeruginosa* followed by *S. aureus* and *A. baumanii* while the highest values were observed for the other Gram-negative rods. However, due to high variability within each species, the differences in sensitivity between species were not statistically significant. There were no significant differences between Gram-negative bacteria and Gram-positive bacteria (in this study represented only by *S. aureus*). It is also apparent that the MIC values for reference strains cannot be considered representative of the clinical isolates. The reference strain of *P. aeruginosa* ATCC 27857 used in the study was less sensitive to AgNPs than *S. aureus* ATCC 29213*, E. coli* ATCC 25922 or *K. pneumoniae* ATCC 31488, while the results obtained for clinical isolates showed a reversed trend.

Antibiotic resistance mechanisms were identified in 14 clinical isolates. Resistance associated with extended-spectrum beta-lactamases (ESBL) was identified in 2 *E. coli* isolates, 2 *K. pneumoniae* isolates and 1 *A. baumanii* isolate. Among the *S. aureus* isolates, 9 isolates were resistant to macrolides, lincosamides and streptogramins (MLS). Two of these isolates were additionally resistant to methicillin (MRSA). Median MIC values for the antibiotic-resistant isolates were higher than for antibiotic-sensitive bacteria, but this difference was not statistically significant (Table 2). However, the number of antibiotic-resistant bacteria in the study was probably too low to observe possible more subtle relationships.

**Table 2.** Minimal inhibitory concentration (MIC) of AgNPs for clinical isolates with and without mechanisms of antibiotic resistance [mg/L]. MIC values did not have a normal distribution, therefore median was used as a measure of central tendency and the interquartile range was used as a measure of statistical dispersion. Higher MIC values indicate lower sensitivity to the AgNPs. ESBL - extended-spectrum beta-lactamases; MRSA - methicillin-resistant *Staphylococcus aureus*; MLS - resistance to macrolides-lincosamides and streptogramins.

| Mechanisms of antibiotic resistance (number of bacterial isolates) | median | min | max | interquartile range |
| --- | --- | --- | --- | --- |
| no mechanisms of antibiotic resistance (34) | 10 | 5 | 40 | 15 |
| with mechanisms of antibiotic resistance (14) | 17.5 | 5 | 40 | 15 |
| ESBL (5) | 25 | 25 | 30 | 2.5 |
| MRSA+MLS (2) | - | 5 | 25 | - |
| MLS (7) | 10 | 10 | 40 | 15 |

The cytotoxicity of AgNPs was assessed in the *in vitro* culture of human primary nasal epithelial cells (HNEpC). Incubation in the presence of AgNPs reduced the viability of epithelial cells in a dose-dependent manner. However, high concentrations of the AgNPs and prolonged incubation time were required to achieve cytotoxic effects (Fig. 3). At 24 hours of incubation, cell viability remained above 80% for up to 75 mg L^−1^ AgNPs, and no morphological changes implicating cell death were observed. At the highest concentration tested (100 mg L^−1^), cell viability measured with the MTT assay was 65.11% +/− 4.86, suggesting that human nasal epithelial cells are much less sensitive to AgNPs than any of the tested bacterial isolates. The half-maximal inhibitory concentration (IC50) for the 48-hour time point was determined at 60.03 mg L^−1^, which was also above the MIC for 100% of the tested pathogens. 80% viability after 48 hours was maintained at 25 mg L^−1^. This concentration of AgNPs was effective against 39 (81%) of the isolates (Fig. 4).

**Fig. 3.**
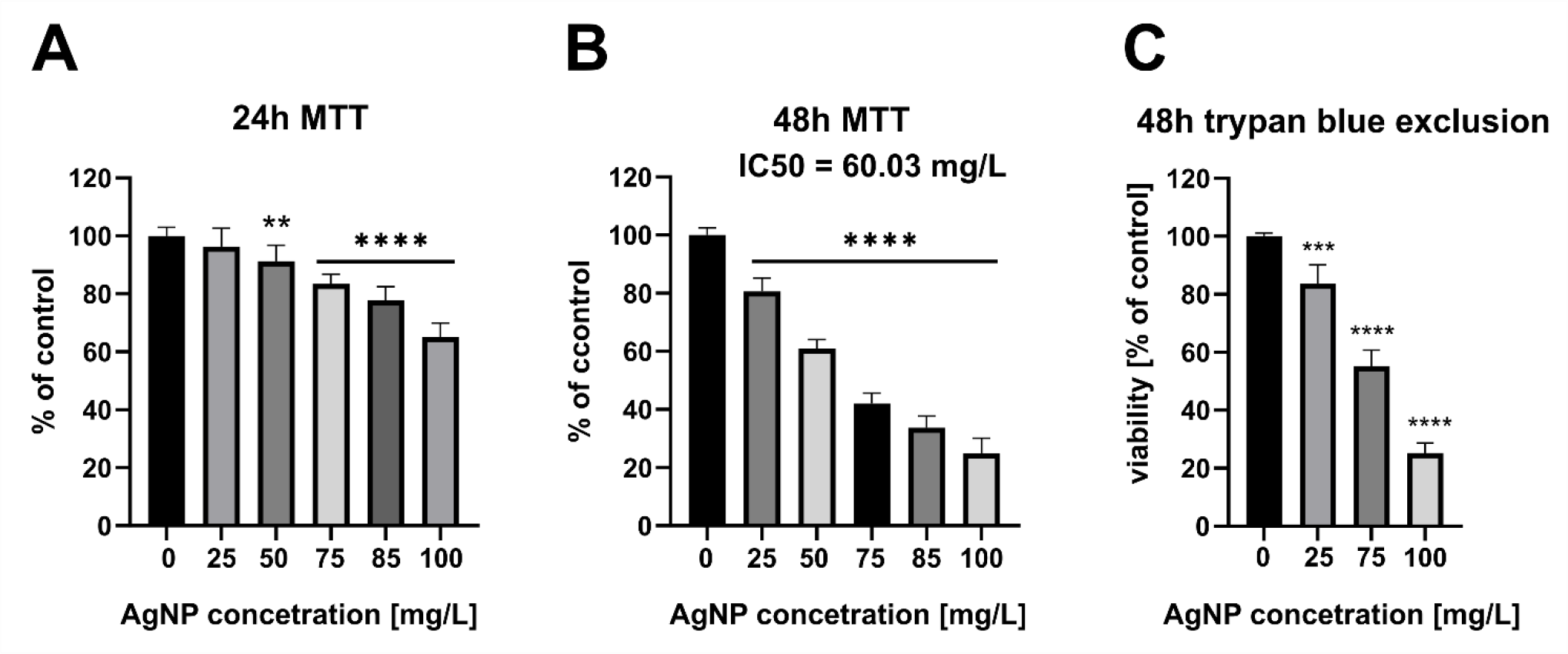
Cell viability of human primary nasal epithelial cells. (A) MTT assay following exposure of cells to AgNPs at the indicated concentrations for 24 hours. (B) MTT assay following exposure of cells to AgNPs at the indicated concentrations for 48 hours. Data represent mean values of three independent experiments performed in triplicate (n=9), expressed as % of control ± SD. * indicates values significantly different from the control i.e., MTT assay for 0 mg L^−1^ of AgNP. (C) Trypan blue exclusion assay. The percentage of viable cells was determined after a 48-hour incubation in the presence of AgNPs. Data represent mean values ± SD. Statistical significance compared to the control assays at 0 mg L^−1^ of AgNP. ** P ≤ 0.01; *** P ≤ 0.001; P **** P ≤ 0.0001.

**Fig. 4.**
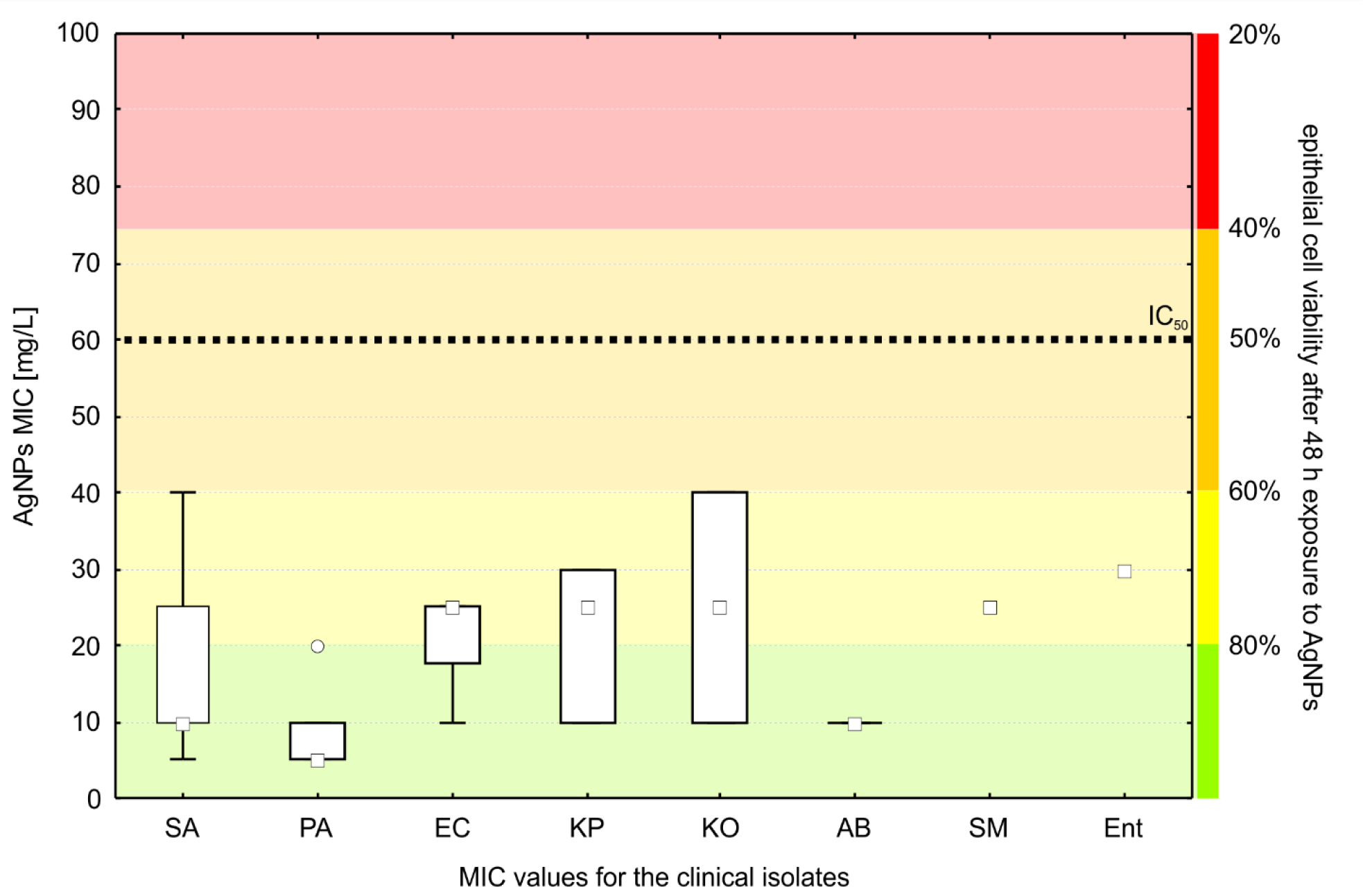
MIC values for the clinical isolates compared to the nasal epithelial cell viability after 48 h exposure to AgNPs (IC_50_ - the half-maximal inhibitory concentration). SA – *Staphylococcus aureus*, PA – *Pseudomonas aeruginosa*, EC – *Escherichia coli*, KP – *Klebsiella pneumoniae*, KO – *Klebsiella oxytoca*, AB – *Acinetobacter baumanii*, SM – *Serratia marcescens*, Ent - *Enterobacter cloacae)*. The boxes represent the interquartile range,

The findings of the colorimetric assay were confirmed with a cell death assay based on cell membrane integrity (trypan blue exclusion assay). Results obtained using both methods were comparable, which indicates that the lower absorbance values observed in the MTT assay are due to the changes in cell viability and not to changes in cell mitochondrial activity or cytostatic effects of the nanoparticles.

## Discussion

In this study, we examined the sensitivity of sinonasal pathogens to AgNPs obtained with the use of tannic acid. The MIC values of the clinical isolates ranged between 5 and 40 mg L^−1^. The values reported by other authors for the same bacterial species vary between studies due to differences in the type of AgNPs and bacterial strains used for sensitivity testing [8, 24–27].

Furthermore, it is a common practice to use only one reference strain to represent a species. Our research proved that this approach is incorrect. We showed that the sensitivity to AgNPs in reference strains was not representative of the values noted for clinical isolates of the same species. The MIC values for the reference strains of *S. aureus E. coli* and *K. pneumoniae* were two to five times lower than the median values for the clinical isolates. In contrast, the reference *P. aeruginosa* was characterized by MIC values that were twice higher than the median value for clinical isolates of the same species.

Among the clinical isolates, there were considerable differences in the MIC values between isolates of the same species. These differences were not species-specific. For most species, we found isolates that were highly sensitive to AgNPs (MIC 5 mg L^−1^) as well as isolates that were inhibited only by much higher concentrations of the antimicrobial (40 mg L^−1^).

Differences in the cell wall structure between Gram-positive and Gram-negative bacteria have been shown to influence their interactions with metal nanoparticles [10]. In several studies, the Gram-negative *E. coli* was more susceptible to AgNPs than the Gram-positive *S. aureus* [28] with 2-10 times lower MIC values depending on the study [24, 25]. On the contrary, in our study, there were no statistically significant differences in MIC values between Gram-positive and Gram-negative bacteria, but the mean MIC value for Gram-positive isolates was lower than for the Gram-negative isolates. The AgNPs used in our study were characterized by negative zeta potential. Therefore, it can be assumed that repulsive electrostatic interactions occur between the AgNPs and the LPS in the cell wall of the Gram-negative bacteria [10]. However, our previous research showed that, apart from electrostatic interactions, multiple other factors, including the stabilizing agents, influence the antibacterial effects of AgNPs [5].

Previously, the efficacy of AgNPs against clinical isolates from patients with CRS and otitis was assessed by Feizi *et al*. [6]. Eighteen bacterial strains were included in the study (5 MRSA, 5 *P. aeruginosa*, 5 *H. influenzae*, 3 *S. pneumoniae*). *P. aeruginosa* was more sensitive to AgNPs than other species, but in general Gram-positive isolates were not universally more resistant than Gram-negative bacteria. The authors also noted that MIC values were different among strains for each of the species. The study was conducted in Australia and utilized different AgNPs than the ones used in our research, but the conclusions are strikingly similar to our observations carried out in Europe. Both stupgdies show that the sensitivity of sinonasal pathogens to AgNPs is not easily predictable. If AgNPs are used in therapy, each isolate obtained from a patient needs to be tested individually to determine possible resistance.

Recent observations suggest that uncontrolled application of silver without prior knowledge about the pathogen’s sensitivity may result in exposition to sublethal concentrations and promote resistance development. Bacteria can develop multiple resistance mechanisms that protect them from silver and AgNPs [9, 10]. It was shown that resistance can be rapidly induced after exposure to increasing concentrations of silver [11] which suggests that silver preparations should be applied with caution. Just like antibiotics, they need to be used only when antimicrobial treatment is clearly indicated, and the dosage must be adequate to achieve concentrations above the MIC values of the targeted pathogen.

Resistance to silver was also shown to promote cross-resistance to antibiotics [11, 29, 30]. In our study, 37.5% of the isolates had mechanisms of antibiotic resistance. We observed higher MIC values for these isolates compared to antibiotic-sensitive bacteria, but the differences did not reach statistical significance.

Studies reporting the prevalence of silver resistance in clinical settings are scarce. However, their results to date seem to be fairly optimistic. Most data on the subject is related to bacteria isolated from burns, ulcers and other wounds that are frequently treated with dressings containing silver. Although silver resistance genes were encountered in 2-6% of clinical isolates, genetic resistance usually did not translate into phenotypic resistance [15, 31–33]. Nevertheless, the first strains that could tolerate silver concentrations up to 5500 uM and were resistant to many commercially available wound dressings were occasionally found in environments where exposition to silver is more frequent [15, 34, 35]. Bacteria that do not display overt Ag-resistance frequently harbour genes that cause cryptic resistance which is readily activated upon silver challenge [36]. These results indicate that silver resistance, although currently rare, may become an increasing problem in the future in case of unlimited overuse of silver preparations.

The question of whether AgNPs can be safely used as antimicrobial drugs in humans is under debate. Silver preparations are available over-the-counter despite a lack of adequate regulatory approval [7]. The toxicity levels reported for eukaryotic cells vary and depend on the properties of the AgNPs used in the experiments and the type of cells under consideration (15 mg L^−1^ for alveolar epithelial cells, 30 mg L^−1^ for monocytes, 80 mg L^−1^ for HeLa epithelial cells) [37–39].

The toxicity of AgNPs for human cells was compared with the toxicity of sinonasal pathogens in two studies. Feizi *et al*. studied 18 clinical isolates from patients with CRS and otitis media. The AgNPs for this study were produced using *Corymbia maculate* leaf extracts. The AgNPs were not toxic to human immortalized bronchial epithelial cells Nuli-1in concentrations equal to MIC for the sinonasal pathogens and up to 175 ppm after 1 hour of exposure [6]. Chen *et al*. studied the antibacterial properties of commercially available 10 nm AgNPs that were effective against reference strains of *E. coli* and *S. aureus* at a concentration of 5 ppm. They found that human nasal squamous cell carcinoma cells (RPMI2650) maintained >80% viability after 24 hours of exposure to 5 ppm of AgNPs [40].

In our study, to assess the cytotoxicity of silver nanoparticles, we used primary HNEpC cells, which are the most reliable two-dimensional *in vitro* model of the nasal epithelium [41]. We observed that the AgNPs were nontoxic to epithelial cells in concentrations that inhibited the growth of all bacterial isolates tested when incubated with the cells for 24 hours. At 48 hours of incubation, silver nanoparticles were not cytotoxic up to 25 mg L^−1^, which is above the median MIC value for the tested bacteria. However, 9 (19%) of the clinical isolates were not sensitive to AgNPs at this concentration.

It is debatable what time of exposure in cytotoxicity experiments is representative of the real-life pharmacokinetics of the AgNPs in the nose and the sinuses. In the study conducted by Feizi *et al*. the incubation time was very short (1 hour). The authors explained that the mucociliary clearance eliminates the AgNPs within about 15 minutes when applied in a nasal solution [6]. However, to fight an infection, it will probably be necessary to maintain the drug concentration above the MIC value for a longer period. For antibiotics, the recommended time of treatment for sinonasal infections is between 5 days and 3 weeks [1]. Therefore, it is reasonable to assume that AgNPs need to be used in a formulation that would increase the retention time of the nanoparticles in the nasal cavity or a dressing that would stay in contact with the nasal mucosa for prolonged periods.

We showed that the AgNPs were not toxic to HNEpC cells at concentrations above MIC for the sinonasal pathogens in the first 24 hours of the experiments, but the cytotoxicity increased when the time of exposure was longer. This observation suggests that the real-life toxicity of AgNPs may be higher than assumed from short-time experiments on cell lines.

On the other hand, several studies indicate that standard two-dimensional *in vitro* cell culture models may overestimate the cytotoxicity of different agents, including silver nanoparticles [42, 43]. Differentiated 3D *in vitro* models might more closely resemble what occurs *in vivo* due to barrier properties that reduce absorption across the stratified epithelium. Zavala *et al*. (2016) showed that higher concentrations and longer exposure times of air pollutants are needed to induce cytotoxic effects in a 3D *in vitro* model of airway epithelium compared to the 2D epithelial cell culture. Chen *et al*. (2019) found no toxicity of AgNPs in a 3D model of a stratified epidermis while observing a pronounced cytotoxic effect of equivalent doses of AgNPs in a 2D culture of keratinocytes. Similar 3D models of differentiated nasal epithelium cultured at the air-liquid interface, containing cilia and mucus-producing cells, have been described [44, 45].

Local toxicity for nasal epithelial cells is not the only factor that limits the potential topical application of AgNPs in patients with CRS. Systemic distribution of AgNPs was observed after intranasal administration in rodent models. Aggregation of AgNPs was noted in the spleen, lung, kidney and nasal airway [14]. This resulted in enhanced destruction of erythrocytes in the spleen, but no other microscopic changes were associated with the AgNPs depositions. It has to be noted that the doses of AgNPs used in the study were exceedingly high (100-500 mg/kg). However, repeated inhalation of AgNPs for 28 days at concentrations of 10^6^ particles/cm^3^ caused no significant changes in the nasal cavity or lungs of rats, only an increase in the number and size of goblet cells. Moreover, direct nose-do-brain transport of AgNPs is suspected to bypass the blood-brain barrier [14, 46, 47], but not all studies support this observation [7]. Importantly, the application of AgNPs results in significantly lower concentrations of silver in blood than the delivery of AgNO_3_ [7].

The safety of AgNP sinonasal rinses (15 mg L^−1^) in 11 patients with CRS was assessed by Ooi *et al*. [12]. Four patients had elevated serum silver levels but no adverse events were reported and the silver levels did not reach the threshold for argyria. In our study, only 27 (56%) pathogens were sensitive to AgNPs at concentrations of 15mg L^−1^. However, Ooi *et al*. applied different AgNPs than the ones used in our study, which excludes any direct comparisons.

In conclusion, the results of our study showed that most pathogens isolated from CRS patients during the episodes of exacerbations were sensitive to AgNPs in concentrations safe for the nasal epithelium *in vitro*. However, *in vitro* experiments may under- or overestimate the toxicity of antimicrobials.

Therefore, further *in vivo* studies on the toxicity, pharmacokinetics and pharmacodynamics are required before AgNPs are approved as intranasal antimicrobials in humans.

Due to the unpredictability and significant differences in the MIC values between isolates, testing of sensitivity to AgNPs should be indicated before their application in every case. This approach may prevent exposition to suboptimal drug concentrations that promotes the development of silver resistance.

## Limitations

(a) The study group included only patients with postoperative AECRS so the results presented in this paper may not apply to bacterial exacerbations of CRS in patients who did not undergo sinus surgery. (b) Only aerobic species were included in the study. (c) In this study, we used one type of AgNPs. The properties of AgNPs obtained with different methods may vary. (d) This observational study was designed to estimate the prevalence of phenotypic resistance while the prevalence of resistance genes or cryptic resistance and determination of resistance mechanisms requires more detailed exploration to explain the observed differences in sensitivity between bacterial isolates. (e) The results of *in vitro* tests of AgNPs efficacy are always affected by the media in which the experiment is conducted due to their interaction with ions, macromolecules and blood [10]. Therefore, further research on the interactions between the AgNPs and the secretions in the sinonasal cavities is necessary before their application *in vivo*.

The authors declare that the research was conducted in the absence of any commercial or financial relationships that could be construed as a potential conflict of interest.

## Acknowledgements

J.S. acknowledges funding from the Jagiellonian University Medical College statutory funds (grant number N41/DBS/000462). A.G., T.G. and M.S. acknowledge funding from InterDokMed POWR.03.02.00-00-I013/16. The funders had no role in study design, data collection and interpretation. The authors acknowledge Bogna Daria Napruszewska for conducting ICP-MS.

## Declarations

### Data availability

The raw data supporting the conclusions of this article will be made available by the authors, without undue reservation; the preprint of this manuscript is available at bioRxiv https://doi.org/10.1101/2022.01.03.474872.

### Ethical approval

This study was granted by the Bioethics Committee of Jagiellonian University Medical College (approval no. 1072.6120.208.2017).

Informed consent was obtained from all individual participants included in the study

## Authors contributions

JSz, JD, MSz, TG - study conceptualization, JSz, AG, JS, MO, PŻ - data curation, JSz, JS - formal analysis, JSz, AG, MO, PŻ – investigation, JS - investigation (cytotoxicity), JSz, TG - funding acquisition, JSz, AG - investigation, JSz, JD, MSz, TG - methodology, JSz, MSz, TG - project administration, JSz, TG - resources, AG - resources (microbiology), JD - resources (cytotoxicity), MO, PŻ - resources (AgNPs), JSz, MSz, TG - supervision, JSz, MSz - visualization, JSz - writing – original draft, JS - writing – original draft (cytotoxicity), JSz, JD, TG, MSz, JS - writing – review and editing

### Consent to participate

Informed consent was obtained from all individual participants included in the study.

### Consent for publication

The authors affirm that human research participants provided informed consent for publication of the data presented in Fig. 2 as well as in Tables 1 and 2.

